# Respiratory shedding of infectious SARS-CoV-2 Omicron XBB.1.41.1 lineage with increased evolutionary rate among captive white-tailed deer

**DOI:** 10.1101/2024.09.24.613938

**Authors:** Francisco C. Ferreira, Tahmina Pervin, Wendy Tang, Joseph Hediger, Logan Thomas, Walter Cook, Michael Cherry, Benjamin W. Neuman, Gabriel L. Hamer, Sarah A. Hamer

**Affiliations:** Texas A&M University, College Station, TX, USA; Texas A&M University-Kingsville, Kingsville, TX, USA

**Keywords:** One Health, whole genome sequencing, COVID-19, *Odocoileus virginianus*, spillover

## Abstract

White-tailed deer (*Odocoileus virginianus*) have high value for research, conservation, agriculture and recreation, and may be important SARS-CoV-2 reservoirs with unknown human health implications. In November 2023, we sampled 15 female deer in a captive facility in central Texas, USA. All individuals had neutralizing antibodies against SARS-CoV-2 and 11 had RT-qPCR-positive respiratory swabs; one also had a positive rectal swab. Six of 11 respiratory swabs yielded infectious virus with replication kinetics of most samples displaying lower growth 24-48 h post infection *in vitro* when compared to Omicron lineages isolated from humans in Texas in the same period. However, virus growth was similar between groups by 72 h, suggesting no strong attenuation of deer-derived virus. All deer viruses clustered in XBB Omicron clade, with more mutations than expected compared to contemporaneous viruses in humans, suggesting that crossing the species barrier to deer was accompanied by a high substitution rate.

## Introduction

The white-tailed deer (*Odocoileus virginianus*, henceforth deer), a native North American species, is highly susceptible to infection with the severe acute respiratory syndrome virus 2 (SARS-CoV-2) (1, 2) and can serve as a viral reservoir as infected free-ranging deer can initiate onward intra-specific transmission (3, 4). SARS-CoV-2 lineages have been detected in deer sampled up to four-five months after the same viral lineage was displaced from human populations by other lineages (5, 6), suggesting sustained transmission within free-ranging animals. Transmission within deer can lead to a gradual accumulation of mutations in animal-adapted SARS-CoV-2 (7), which could facilitate immune evasion when transmitted back into human populations (8). Additionally, detections of deer-adapted virus infecting humans in Canada (9) and in the US (3) raise concerns about the emergence of sustained human-deer-human SARS-CoV-2 transmission cycles.

Captive animal herds with increased population densities and frequent human intervention may support transmission of zoonotic pathogens (10, 11). Captive deer were also found to be naturally exposed to SARS-CoV-2, as demonstrated by high rates of detection of viral RNA (12) and of neutralizing antibodies at high titers among deer (13). Managing highly susceptible animal species under human care creates opportunities for human-to-deer transmission with subsequent rapid dissemination within herds (14). Data collected in 2012 estimated that there are around 5,500 white-tailed deer breeding facilities in the US (15). In Texas for instance, the more than 1000 existing facilities manage an average of 180 to 242 deer each (16, 17). Consequently, such large numbers of deer under human management may favor human-to-deer and deer-to-deer transmission events with subsequent spread of deer-adapted viruses from farmed animals back into humans, which has been documented with farmed minks (*Neovison vison*) in Europe (18, 19) and possibly in the USA (20).

Despite continued testing of deer for SARS-CoV-2 since the Omicron variant of concern (VOC) emerged in the US in November 2021 (e.g., (3, 21)), the detection rate of recent lineages in deer remains low. For example, only 2.5% (16 out of 648 genomes) of the deer-derived sequences belong to the Omicron clade (GRA) as per GISAID (the Global Initiative on Sharing All Influenza Data; (22)) database (accessed on August 14^th^, 2024). These findings suggest deer are susceptible to new emerging lineages, yet it is unclear whether a salient reduction in infection is due to reduced deer exposure to SARS-CoV-2, reduced virus fitness and detectability, reduced surveillance efforts, a lag between detection and reporting, or a combination of factors. Therefore, continued genomic surveillance of highly susceptible species remains critical from a One Health perspective to understand the current landscape of SARS-CoV-2 transmission and evolution in wildlife reservoirs.

We established an active surveillance program to determine SARS-CoV-2 circulation in captive deer in Texas to understand and mitigate transmission risk between humans and animals. Here, we describe a SARS-CoV-2 outbreak in a captive deer breeding facility in November 2023.

## Methods

### Cervid facility

On November 15, 2023, we visited a private cervid facility in Milam County, Texas. Deer were restricted to two pens separated by a fence. This facility falls into the category of “outdoor ranch” as described by Rosenblatt et al. (23), although human-deer contact rates are high in our case due to veterinary care and animal husbandry.

### Deer sampling

Fifteen apparently healthy female white-tailed deer were chemically immobilized for veterinary care at which time samples were collected for the SARS-CoV-2 investigation. We collected oral and nasal swabs from deer using sterile polyester-tipped applicators with polystyrene handles (Puritan Medical Products, Guilford, ME, USA) and combined (henceforth respiratory swab) for each animal into a vial containing 3 mL of viral transport media (VTM; made following CDC SOP#: DSR-052-02). We collected rectal swabs from the same individuals into a separate vial with VTM. We collected blood (approximately 10 mL) from all individuals via jugular venipuncture using sterile needles and syringes, transferred to tubes without anticoagulants and aliquoted the obtained serum samples into microtubes. Our study followed relevant guidelines and regulations approved by the TAMU’s Institutional Animal Care and Use Committee and Clinical Research Review Committee (2022-0001 CA).

### Molecular detection and whole genome sequencing

We extracted total nucleic acid (TNA) from respiratory and rectal swabs using the MagMax CORE Nucleic Acid Purification Kits on a 96-well Kingfisher Flex System (ThermoFisher Scientific, Waltham, MA, USA) following manufacturer’s instructions using 200 μL of VTM from each sample, and 90 μL of elution buffer. We added phosphate buffered saline 1× and a plasmid containing a portion of the RNA-dependent RNA polymerase SARS-CoV-2 gene to the TNA extraction plate as negative and positive extraction controls, respectively. We tested 5 μL of TNA for SARS-CoV-2 by RT-qPCR targeting the RdRp gene of the virus following reagent concentrations as described in Corman et al. (24). We utilized negative and positive controls as used for TNA extraction also for RT-qPCR assays and yielded their expected results. We considered samples with cycle threshold values < 40 as positive.

We submitted all RT-qPCR-positive samples to whole genome sequencing at the Texas A&M University TIGSS Molecular Genomics Core. For each sample, 20 μL of TNA was submitted to library preparation using the xGen SARS-CoV-2 Amplicon Panel (Integrated DNA Technologies -IDT). Sequencing was performed in an Illumina NovaSeq SP PE 2 × 150 flowcell v1.5 to generate an average of 3 million reads per sample, which were initially mapped and assembled using the Illumina DRAGEN-covid-pipeline-RUO-1.0.0.

We interrogated positions with missing sequence data or mixed sequence data, indicated by IUPAC nucleotide uncertainty codes K, S, R, Y, M, W or N in the assembled genome by BLAST 2.15.0, to verify the presence of indels and generate a majority-rule consensus sequence for each sample, which was used in subsequent phylogenetic analysis. We analyzed consensus sequences on Nextclade v3.2.0 (https://clades.nextstrain.org) for clade and lineage assignment (25).

### Bioinformatics

We investigated Mutations, insertions and deletions in the consensus sequences using NextClade and the Global Initiative on Sharing All Influenza Data (GISAID dataset; Shu and McCauley (22)). Sequence differences are reported relative to the genetically closest strain to the deer-derived SARS-CoV-2 isolates, hCoV-19/USA/CA-HLX-STM-DTHMADVDZ/2023, collected from a human in California on November 11, 2023. We used UShER (https://genome.ucsc.edu/cgi-bin/hgPhyloPlace; (26), assessed August 14, 2024) to determine the 10 SARS-CoV-2 available genomes most closely to the sequences obtained in this study.

### Virus isolation

We isolated SARS-CoV-2 in a biosafety level 3 laboratory at the Global Health Research Complex (GHRC) at Texas A&M University by passaging a mix of 100 μL of VTM with 900 μL of 1× Dulbecco’s Modified Eagle Medium (DMEM) via syringe filtration using an 0.2 micron pore size onto onto Vero E6-TMPRSS2-T2A-ACE2 (BEI Resources, NR-54970) cells expressing both endogenous cercopithecine ACE2 and TMPRSS2 as well as transgenic human ACE2 and TMPRSS2. We cultured these cells in DMEM supplemented with 10% fetal bovine serum, 1× antibiotic/antimycotic and puromycin dihydrochloride (10 µg/ml final concentration). Cell plates were incubated at 37ºC with 5% CO_2_, and the supernatants from samples exhibiting cytopathic effects within 24-72 h were collected for titration of infectivity.

We performed titration of infectious virus by using the tissue culture infectious dose 50% (TCID_50_) method on onto Vero E6-TMPRSS2-T2A-ACE2 cells. Briefly, we prepared serial three-fold dilutions of inocula in culture medium and then used them to inoculate monolayers of cells in tissue culture-treated 96-well plates. After 72 h, we fixed the cells with phosphate buffered physiological saline containing 25% final concentration of formalin and stained with crystal violet. We calculated titers of infectious virus using the Reed-Muench method (27).

### Virus growth curves

We used freshly-titrated virus stocks with equal infectivity to inoculate onto Vero E6-TMPRSS2-T2A-ACE2 cells for 30 min, removed inoculum by three rinses with phosphate-buffered saline, and replaced with culture medium. We obtained through GHRC contemporaneous human clinical samples from nasal swabs that were collected at various medical centers in southeastern Texas as part of an ongoing Texas Department of State Health Services virus surveillance project. We worked under approval from the Texas A&M University Institutional Review Board. The samples selected for use here included one isolate from each Pango lineage (28, 29) obtained from patient clinical samples at our coronavirus disease 2019 (COVID-19) screening facility between October 1 and November 30, 2023, namely Pango lineages HV.1, HY.1, HK.11, HK.3, EG.10, EG.5 and JD.1 and were processed (isolated and titrated) in parallel with deer-isolated viruses. These human-derived isolates include all the distinct Pango lineages that were obtained one month before to one month after the deer-derived samples. Additionally, the selected lineages represent the genotypes of 864 of 2253 total (38%) sequenced SARS-CoV-2 complete genomes (excluding low-coverage, and partial sequences) reported to the GISAID from Texas between October 1 and November 30, 2023. Performing experiments in parallel using human and deer samples enabled us to compare *in vitro* growth rates for both sample types. Small aliquots (100 μL) of medium were collected at eight, 24, 48 and 72 h after inoculation, submitted to RNA extraction and submitted to a Luna One-Step RT-qPCR (NEB, Ipswich, MA, US) targeting the nucleoprotein gene. We conducted a One-Way ANOVA with Tukey’s correction for multiple comparisons to compare growth curves of human- and deer-derived isolates. Growth curve plots were generated using GraphPad (version 10.0.0).

### Plaque reduction neutralization tests

Sera from all deer were tested for the presence of neutralizing antibodies against SARS-CoV-2 (isolate USAIL1/2020, NR 52381; BEI Resources, Manassas, VA, USA) via plaque reduction neutralization tests (PRNT) in Vero-CCL-81 (13). Aliquots of the samples were inactivated at 56ºC for 30 min and screened at an initial dilution of 1:10. Samples that neutralized at least 90% of viral plaques, in comparison to the virus control, were further tested with serial 2-fold dilutions to determine the 90% endpoint titers (PRNT_90_).

## Results

### SARS-CoV-2 detection and serology in white-tailed deer

The respiratory swabs of 11 of the 15 deer (73.3%) tested positive for SARS-CoV-2; one of these positive deer also had a positive rectal swab (D89) whereas all other rectal swabs were negative. The mean for the Ct values for respiratory samples was 27.93 (SD = 3.20) ranging between 22.12 and 31.92.

All 15 deer had neutralizing antibodies against SARS-CoV-2 as demonstrated by PRNT. PRNT_90_ endpoint titers varied from 1:10 to 1:320 in RT-qPCR-positive animals (mean 90; SD = 92) and ranged from 1:20 to 1:640 in RT-qPCR-negative animals (mean 190, SD = 301).

### Virus isolation

To assess whether deer-derived SARS-CoV-2 was able to efficiently use human ACE2 and TMPRSS2 for cell entry, we inoculated Vero E6-TMPRSS2-T2A-ACE2 cells with all 12 RT-qPCR-positive respiratory and rectal swab samples. Six respiratory samples (D92, D93, D94, D95, D96, D100, Figure 1) produced cytopathic effects characteristic of SARS-CoV-2 on this cell line, including extensive syncytium formation followed by rounding and detachment, and each of these viral isolates was designated as “recovered” to differentiate the isolates from the potentially broader SARS-CoV-2 populations present in samples from infected animals. Stocks of each isolate were grown and titrated by the TCID_50_ method for further study.

**Figure 1.**
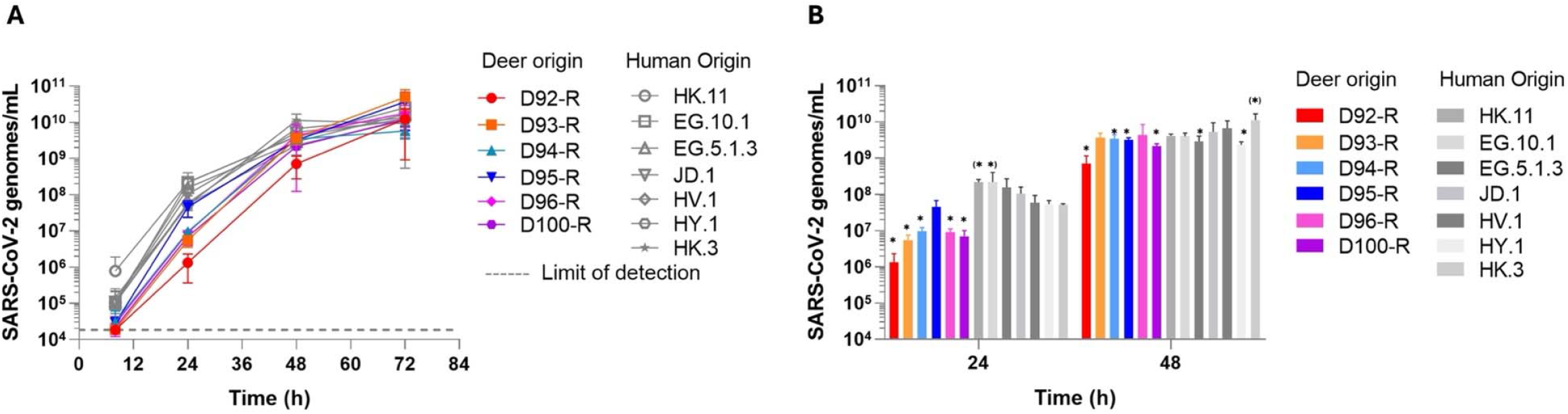
Multi-step growth characteristics of contemporaneous strains isolated from captive white-tailed deer and humans. (A) Vero E6-TMPRSS2-T2A-ACE2 cells were inoculated with SARS-CoV-2 recovered from deer (D92-R to D100-R) or from human clinical nasopharyngeal samples (EG.10.1, EG.5.1.3, HK.11, JD.1, HV.1, HY.1, HK.3) at a multiplicity of infection of 0.002. Samples of the supernatant were collected and titrated by one-step quantitative PCR. Error bars show the standard deviation calculated from three replicates. The lower limit of detection is indicated with a dashed line. Averaged data for the 24 h and 48 h timepoints are shown in panel B. Statistically significant differences after one-way ANOVA with Tukey’s correction are indicated by asterisks (for values significantly lower than aggregated human samples) and asterisks in parentheses (for values significantly higher than aggregated human samples). We added “-R” to the name of each animal ID to indicate that samples used in these experiments were recovered from the initial virus isolation step.

To compare growth rates of viruses isolated from deer with strains that were circulating in humans in Texas at the same time, we performed multi-step growth curves, starting at low multiplicity of infection (0.002). Five and four out of six deer-derived viruses showed lower growth rates at 24 h and 48 h, respectively, when compared to human-derived isolates (Figure 1; *P* < 0.05). There were no differences in growth at 8 h and at 72 h (Figure 1). This data indicated that deer-derived SARS-CoV-2 isolates were able to efficiently grow in cells expressing human ACE2 and TMPRSS2, with a delay in early growth and no difference in burst size compared to contemporaneous SARS-CoV-2 isolated from humans.

### SARS-CoV-2 sequencing

We obtained complete (>99.8%) SARS-CoV-2 genomes from all 12 RT-qPCR-positive samples (11 respiratory and one rectal swab) belonging to 11 animals. Analysis by Nextclade revealed that all viruses fall within the XBB clade (XBB.1.41.1 lineage assigned by Pangolin version v4.3, also known as JC.5). Viruses most closely related to the ones sequenced from deer were collected from humans in California, November 11, 2023 (GISAID accession hCoV-19/USA/CA-HLX-STM-DTHMADVDZ/2023), from Texas December 19, 2023 and several other strains from Texas in January, 2024 (Figure 2). These sequences from humans were obtained from different GISAID submitters and had identities ranging between 95.5% (1,352 bases different) and 99.9% (21 nucleotides different) in relation to the sequences we obtained from deer. No SARS-CoV-2 detected in deer in other regions of North America clustered together with the genomes obtained here.

**Figure 2.**
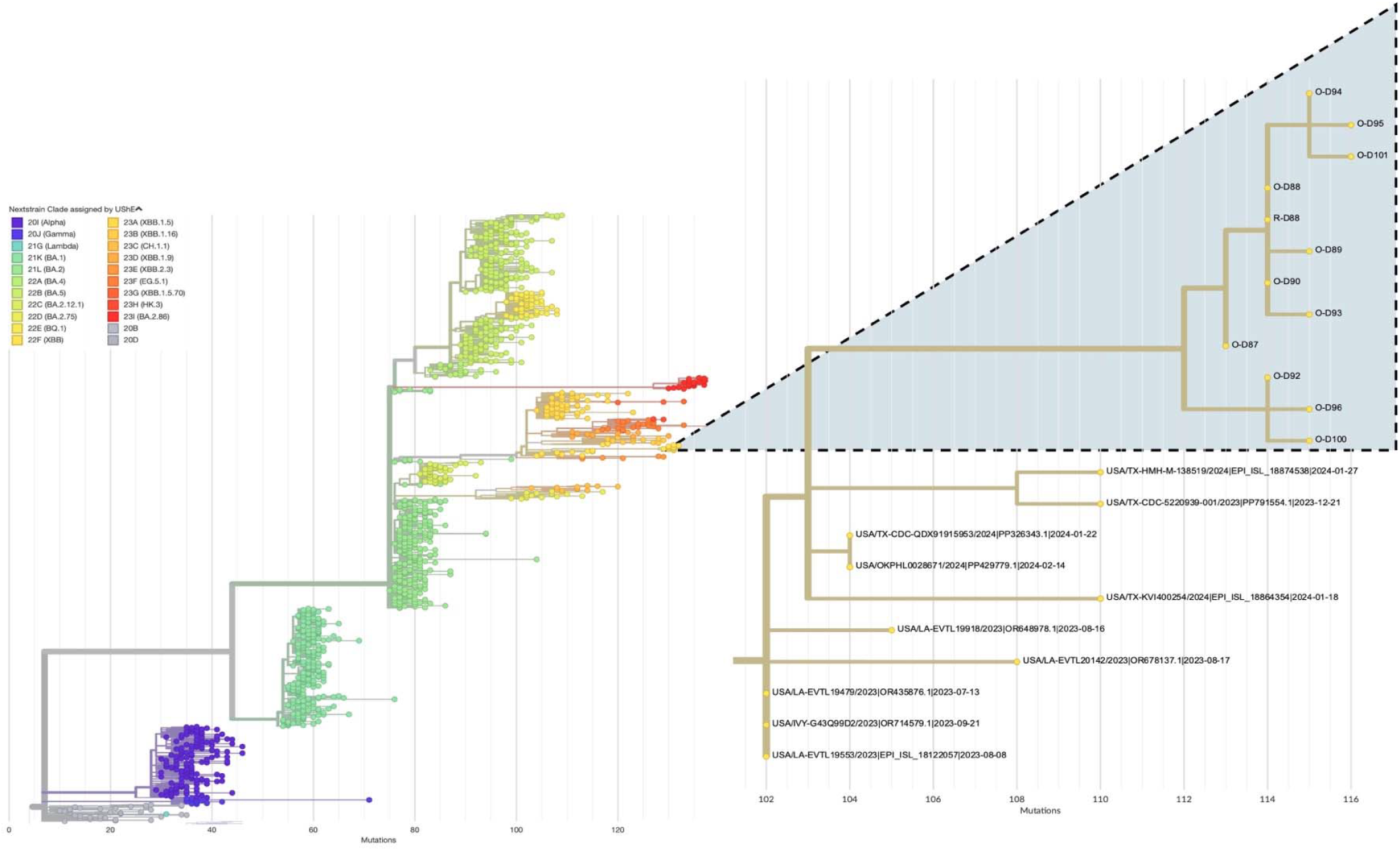
Phylogenetic context of SARS-CoV-2 from captive white-tailed deer sampled in Milam County, Texas, November 2023. We analyzed genome sequences from deer with genomes representative of the main virus clades in the main phylogenetic tree. The secondary tree displays the placement of genomes obtained in our study (highlighted clade; O = oral/nasal swab; R = rectal swab) in relation to 10 of the most closely related samples deposited in GISAID as of August 14, 2024.

Our deer-derived sequences had 15 nucleotide substitutions not found in the closest human-derived isolates and up to 11 substitutions shared by some or none of the other deer-derived sequences. Compared to these closely related human samples, the viruses detected in deer did not have unique mutations in the spike (S) protein. However, the viruses detected from humans in Lubbock Co. and in Oklahoma had amino acid changes (N74S and Q146H, respectively) not present in the deer samples when using the lineage XBB as a reference.

## Discussion

All 15 deer sampled in a captive facility in Texas had evidence of exposure to SARS-CoV-2, with 11 (73%) of them shedding viral RNA, showing a widespread virus dissemination among these animals. Additionally, whole genome sequencing revealed recent Omicron lineages continue to be detected within deer in North America. From these RT-qPCR-positive deer assessed, six had infectious virus in their upper respiratory tract. Although viruses isolated from most deer showed slower growth at 24 h and 48 h post inoculation *in vitro* when compared to contemporaneous human-isolated viruses, the number of viral copies at 72 h was similar between groups, showing no attenuation in deer-adapted viruses. The XBB lineage detected in our study was circulating at low frequency in human populations in the United States, with a cumulative prevalence lower than 0.5% between May 2023 and January 2024 (30), concurrent with the period when deer were sampled for this study.

Human-derived SARS-CoV-2 have 30-32 substitutions per genome per year (around 2.5/month; https://nextstrain.org/ncov/gisaid/global/6m?l=clock; accessed on August 13, 2024). The human strain hCoV-19/USA/CA-HLX-STM-DTHMADVDZ/2023 and our deer samples were collected seven days apart, and displayed 15-22 nucleotide substitutions among them, suggesting that crossing the species barrier from human to these captive deer was accompanied by a high substitution rate. However, it is difficult to determine this pace without knowing when the initial spillover into this population occurred. Possible explanations for this proposed burst of evolution include bottlenecking and strong activation of mutagenic intracellular factors such as RNA editing enzymes (ADAR1) and apolipoprotein B mRNA editing enzyme (APOBEC) in deer (9). For instance, the latter innate immune response factor play a key role during natural infections, and the catalytic subunit 3H (APOBEC3H) was shown to display stronger upregulation in infected deer when compared to natural infections in humans (31). Furthermore, the comparable to slightly attenuated growth of deer and human isolates suggests that deer-to-human transmission, and concomitant bursts of evolution within deer, have the potential to produce variants that may not display substantial loss of adaptation for infection and development in humans. This suggests that Omicron SARS-CoV-2 lineages circulating in deer may be efficiently transmitted back to humans.

The XBB clade has been detected in multiple wildlife species in areas within a rural-to-urban gradient, with at least five transmission events from human to animals (32). Because wildlife species susceptible to SARS-CoV-2 share habitat with captive deer (32, 33), our results suggest that infectious deer have the potential to infect and be infected by commingling species. Assessments of wildlife, livestock and humans that commingle with infected captive deer are a high priority for testing for cross-species transmission events.

Free-ranging deer are predicted to be at higher risk of exposure to SARS-CoV-2 when sharing fence line with captive populations (23), and the high frequency of infectious deer detected here emphasizes that captive-to-wild transmission between deer should be empirically tested. Additionally, fence line transmission could explain, at least partially, the widespread sustained SARS-CoV-2 transmission within wild deer in North America (34).

Some wildlife species found shedding RNA of XBB lineages under natural conditions (e.g.; raccoons -*Procyon lotor*, and cottontail rabbit -*Sylvilagus floridanus*; (32)) did not show clinical signs of infection, nor did shed viral RNA when experimentally infected with early, pre-Omicron lineages (35, 36). These studies together provide strong evidence that, as new lineages emerge, the breadth of competent hosts for the virus changes, highlighting the importance of continued surveillance to assess risks of establishment of transmission cycles of SARS-CoV-2 among wildlife species.

Infectious virus can be isolated from nasal swabs up to 6 days post-infection in deer, while viral RNA can be detected in respiratory swabs up to 22 and 21 days post direct experimental infection and post contact with infected deer, respectively (1, 2). These studies, which used pre-Omicron lineages revealed efficient horizontal transmission among deer. Six out 11 RT-qPCR-positive deer from out study had infectious virus, suggesting they became infected within about six days before our sampling. Our results show that more recent lineages are also efficiently and quickly transmitted amongst deer. However, controlled laboratory experiments are needed to confirm the duration of infectious virus shedding during infections with more recent SARS-CoV-2 lineages in deer.

Our study demonstrates the continued and efficient spread of SARS-CoV-2 among deer as new virus lineages emerge and infect non-human species, with rapid spread within deer of a XBB lineage concurrently circulating in human populations. These findings corroborate epidemiological models predicting that deer facilities favor high virus dissemination within captive populations (23), and transmission potential may vary across the spectrum of scenarios in which deer and humans interact (e.g. deer ranches, wildlife sanctuaries, safaris, zoos, outdoor recreation with free-ranging animals). Given the uncertain public health and animal health implications of viral maintenance within captive deer, agricultural biosecurity practices may be useful in reducing the possibility of establishing long-term animal reservoirs for the virus. This is important because the mid-long-term evolutionary consequences for SARS-CoV-2 circulating in non-human populations is unknown, and captive facilities may provide opportunities for sustained transmission at this human-deer-wildlife interface. Our study cannot determine how SARS-CoV-2 entered the target deer population. This is an important gap that needs to be addressed to mitigate transmission risks among humans, deer and other commingling species.

## Acknowledgements

Funding was provided by the American Rescue Plan Act through USDA APHIS A1181 Agricultural Biosecurity 2023-70432-39559 and by Texas A&M AgriLife Research. Use of the Global Health Research Complex and the Texas A&M Molecular Genomics Core is acknowledged. The following reagent was obtained through BEI Resources, NIAID, NIH: *Cercopithecus aethiops* Kidney Epithelial Cells Expressing Transmembrane Protease, Serine 2 and Human Angiotensin-Converting Enzyme 2 (Vero E6-TMPRSS2-T2A-ACE2), NR-54970. We thank deer producers across Texas. Laboratory support was provided by Sankar Chaki and Lisa Auckland. Tom Gary DVM, Isabel McAllister and Yuexun Tian assisted in the field. The findings and conclusions in this publication are those of the authors and should not be construed to represent any official USDA or U.S. Government determination or policy.

## Data availability

Virus genome sequences are available in GISAID under Accession numbers EPI_ISL_19411338-48, EPI_ISL_19413822. Raw sequencing data is available in NCBI (BioProject PRJNA1149050).

## Biographical sketch

Dr. Francisco C. Ferreira is an Assistant Research Scientist in the Entomology Department at Texas A&M University. He is interested in wildlife and zoonotic pathogens at the human-animal-environment interface.

